# Conserving a threatened North American walnut: a chromosome-scale reference genome for butternut (*Juglans cinerea*)

**DOI:** 10.1101/2023.05.12.539246

**Authors:** Cristopher R. Guzman-Torres, Emily Trybulec, Hannah LeVasseur, Harshita Akella, Maurice Amee, Emily Strickland, Nicole Pauloski, Martin Williams, Jeanne Romero-Severson, Sean Hoban, Keith Woeste, Carolyn C. Pike, Karl C. Fetter, Cynthia N. Webster, Michelle L. Neitzey, Rachel J. O’Neill, Jill L. Wegrzyn

## Abstract

With the advent of affordable and more accurate third generation sequencing technologies and the associated bioinformatic tools, it is now possible to sequence, assemble, and annotate more species of conservation concern than ever before. *Juglans cinerea*, commonly known as butternut or white walnut, is a member of the walnut family, native to the Eastern United States and Southeastern Canada. The species is currently listed as Endangered on the IUCN Red List due to decline from an invasive fungus known as *Ophiognomonia clavigignenti-juglandacearum* (Oc-j) that causes butternut canker. Oc-j creates visible sores on the trunks of the tree which essentially starves and slowly kills the tree. Natural resistance to this pathogen is rare. Conserving butternut is of utmost priority due to its critical ecosystem role and cultural significance. As part of an integrated undergraduate and graduate student training program in biodiversity and conservation genomics, the first reference genome for *Juglans cinerea* is described here. This chromosome-scale 539 Mb assembly was generated from over 100X coverage of Oxford Nanopore long reads and scaffolded with the *Juglans mandshurica* genome. Scaffolding with a closely related species oriented and ordered the sequences in a manner more representative of the structure of the genome without altering the sequence. Comparisons with sequenced Juglandaceae revealed high levels of synteny and further supported *J. cinerea’s* recent phylogenetic placement. Comparative assessment of gene family evolution revealed a significant number of contracting families, including several associated with biotic stress response.

## Introduction

Temporal analysis of biodiversity loss indicates that we are in the midst of the Earth’s sixth mass extinction event (Ceballos et al. 2017). Habitat destruction and degradation, overexploitation, pollution, invasive species, climate change, and other anthropogenic factors are the greatest threats to Earth’s flora and fauna. The International Union for the Conservation of Nature (IUCN) Red List noted that nearly 30% of the species assessed are at risk of extinction (IUCN 2022), including about 30% of trees.

Preserving adaptive potential through conservation genetics is widely recognized as important to ensuring long-term survival, especially in the face of climate change (Kardos et al 2021). Conservation managers and geneticists can work together to monitor population genetic diversity and develop restoration plans including assisted breeding. However, several studies have acknowledged a “conservation genetics gap” stemming from, among other things, a lack of training, barriers to collaboration, discipline-specific terminology, and access to resources (Taylor et al. 2017; Sandstrom et al. 2019). The slow advancement of tool development and implementation in conservation is further impacted by the lack of genomic resources for the vast majority of threatened species, with less than 1% of all eukaryotic species sequenced to date (Lewin et al. 2022). As such, the implementation of genetic tools in conservation and/or restoration benefits from the development of a reference genome (DeWoody et al. 2022).

Butternut (*Juglans cinerea* L.) is a deciduous, outcrossing, and wind-pollinated canopy tree native to eastern North America. Once valued for its wood and edible nuts, today, it is threatened by butternut canker, a disease caused by the fungus *Ophiognomonia clavigignenti-juglandacearum* (Nair et al. 1979), discovered in the late 1960s and now found throughout butternut’s range. The disease had already killed more than 80% of butternut trees in numerous states by the 1990s (the last detailed censuses available) and is threatening the species with extinction. Diseased trees typically die within 5 to 10 years, and little resistance has been identified outside of hybrids with the introduced Japanese walnut (*Juglans ailantifolia Carrière*). The search for butternut individuals or populations with less susceptibility to butternut canker is confounded by environmental factors. For example, the severity of butternut canker as well as Armillaria root rot is greater in low-elevation sites with heavy soils and in humid sites with heavy canopies (LaBonte et al. 2015). Dramatic declines over the past few decades have resulted in the listing of butternut as endangered in Canada (Nielsen et al. 2003) and Endangered on the IUCN Red list in 2018. Several US states have also included butternut as protected or endangered, including Minnesota and Wisconsin (Smith 2018; WDNR 2021).

Butternut shows high genetic diversity (moderate cline from south to north) (Hoban et al 2008; 2010). In more isolated populations, lower genetic diversity and inbreeding are observed. Chloroplast markers have revealed population-level genetic differentiation across an east–west gradient that may have originated from multiple Ice Age refugia (Laricchia et al 2015). Butternut is sympatric with *Juglans nigra Linnaeus* through much of its range however it cannot hybridize with this species, but it easily hybridizes with its close relative, the non-native *J. ailantifolia.* Hybrids with *J. ailantifolia* are more tolerant of butternut canker, grow faster/larger, and are more productive (Boraks and Broders 2014; Brennan et al. 2020). Hybrids between the species occur at low frequencies in numerous forests in several states (Hoban et al. 2012). Given the limited resistance observed among pure *J. cinerea* individuals, a plan proposed by practitioners seeks to conserve native germplasm and introduce genes for resistance from *J. ailantifolia* (Pike et al. 2020), similar to efforts in American chestnut (Fernandes et al. 2022). The development of reliable markers and the materials for forward and/or reverse breeding approaches, and for analyzing adaptive introgression of hybrids in the wild, will rely on a high quality reference genome.

To address the conservation genetics gap, undergraduate and graduate students trained in the newly formed Biodiversity and Conservation Genomics Center at the University of Connecticut worked with practitioners to generate the first reference genome for the threatened *J. cinerea*. This study represents the intersection of industry (Oxford Nanopore), federal research agencies (USDA Forest Service, Canadian Forest Service (NRCAN)), forest tree conservation (Morton Arboretum), and academia. The chromosome-scale reference was annotated with existing transcriptomic (RNA-Seq) resources and examined in an evolutionary context to recently sequenced members of the Juglandaceae from around the globe.

## Materials & Methods

### Plant Material, Extraction & Sequencing

Leaves were collected from a single individual from a wild population of trees in Sheffield, New Brunswick, Canada (45.88 N, -66.36 W) (Fig. 1A) that appeared healthy (Fig. 1B). A vouchered specimen was deposited at the Connell Memorial Herbarium, University of New Brunswick (UNB #69382). Leaves were flash frozen in liquid nitrogen, and stored at -80°C until extraction. Leaf tissue weighing 750mg was bulked from a single accession and ground with a mortar and pestle in liquid nitrogen. High molecular weight (HMW) genomic DNA extraction was then conducted following Vaillancourt and Buell CTAB-Genomic-tip protocol (Vaillancourt et al. 2019). In brief, the aqueous layer containing the DNA was collected and mixed with isopropanol to precipitate the DNA, which was then resuspended in Buffer G2 and purified using a Qiagen Genomic-tip column. After being eluted in Buffer QF, the purified DNA was further purified by precipitation with isopropanol, followed by a wash with ethanol and elution in Buffer EB overnight at room temperature. DNA was stored at 4°C until shipped to University of Connecticut for library preparation and sequencing. The DNA was sheared to approximately 25 kb using a Covaris^®^ G-tube (#520079, Covaris, LLC, Woburn, MA, USA). The sequencing library was prepared using the ligation sequencing DNA v14 method (SQK-LSK114, Oxford Nanopore Technologies, Oxford, UK). Three separate loads of the library were run on one flow cell (FLO-PRO114M) over 90 hours, resulting in 5.2 million reads with MinKNOW^®^ (v. 22.08.6).

**Figure 1.**
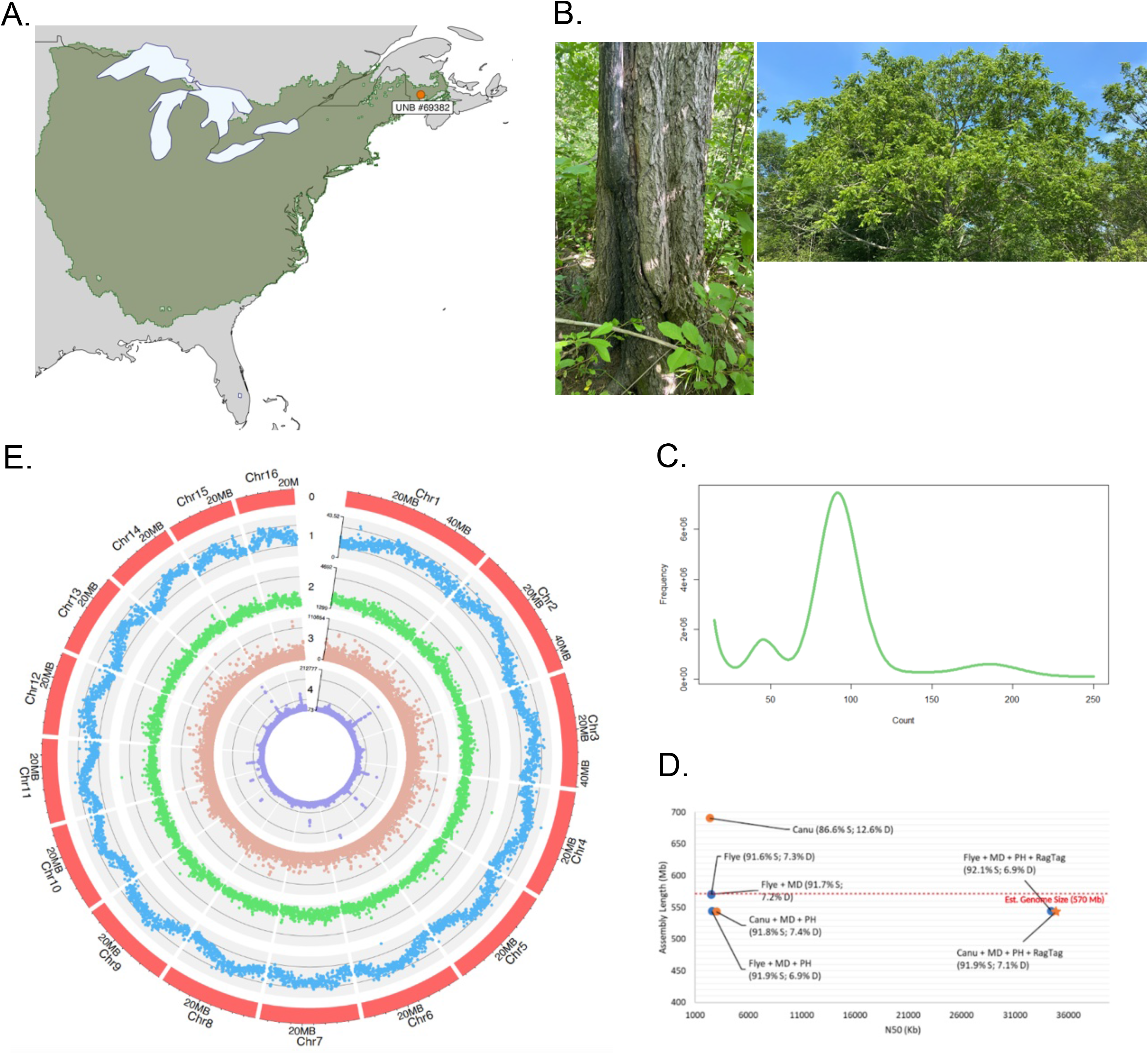
Range map and sample location of the Juglans cinerea reference tree (UNB #69382). Range information from the BIEN database **(A)**. Photographs of the sampled reference tree from the Sheffield, NB population exhibiting a healthy crown and trunk absent of cankers **(B)**. k-mer plot (length=17bp) of k-mer count (x-axis) versus frequency (y-axis). Genome size from long-read data initially estimated at 570 Mb **(C)**. Scatter plot comparing assembly length (Mb) and N50 (kb) obtained from QUAST for Flye and Canu throughout the assembly process. Assemblies represented by color (Canu=yellow; Flye=blue) and shape (drafts=circles; selected final assembly pre-chromosome filtering=star) with abbreviated assembly software (MD=Medaka; PH=Purge Haplotigs). BUSCO single (S) and duplicated (D) completeness scores in parentheses **(D)**. Circos plot representing 0) assembly chromosomes 1) average 5mC methylation in all sequence contexts (CG, CHG, and CHH) over 100 kb windows 2) CpG site count over 100 kb windows 3) gene coverage over 10 kb windows 4) repeat coverage over 10 kb windows **(E)**.

### Assembly

Kmer-freq and GCE were run to estimate genome size at k-mer size 17, taking the long reads (FASTQ) as input (Wang et al. 2020; Liu et al. 2013). A haploid chromosome-scale assembly was generated using Oxford Nanopore PromethION long reads at 100X coverage, based on the estimated genome size of 570 Mb (Fig. 1C; Fig. S1). The raw reads were basecalled by Guppy (v.6.3.8) in two modes: (1) high accuracy (Q>9) duplex (HAC) and (2) super accuracy (Q>10) simplex (SUP). Both sets were independently assessed through the first stages of QC and assembly. To remove potential contaminants from both read sets, including bacteria and fungal associations, Centrifuge (v.1.0.4-beta) was used (Table S1) (Kim et al. 2016). Quality assessment was performed with NanoPlot before and after contaminant removal (De Coster et al. 2018). Two long-read assembly methods, Flye (v.2.9) and Canu (v.2.2), were evaluated (Kolmogorov et al. 2019; Koren et al. 2017). Following *de novo* assembly, Medaka (v.1.7.2) was used for error correcting (polishing) the Flye-generated assemblies (*medaka*). Polishing was not independently run on the Canu assembly outside of Canu’s approach. Completeness and contiguity were assessed at each stage with BUSCO (v.5.0.0) against the Embryophyta single-copy ortholog database (OrthoDB v.10.0), QUAST (v.5.0.2), Minimap2 (v.2.15), and PycoQC (v.2.5.2) (Manni et al. 2021; Kriventseva et al. 2019; Gurevich et al. 2013; Li 2018; Leger and Leonard 2019). These metrics led to the selection of the SUP assemblies for further refinement. Following error-correction, Purge Haplotigs (v.1.1.2) was applied to the Canu and Flye assemblies to reduce allele-based duplications (Roach et al. 2018). Both assemblies were subsequently scaffolded with the chromosome-scale *Juglans mandshurica* reference (BioProject: PRJCA006358) using RagTag (v.2.1.0) in *scaffold mode* (Li et al. 2022b; Alonge and Leonard 2021). The assemblies were then filtered to include the 16 longest scaffolds that aligned to the 16 chromosomes. Using the same assessments listed above, the scaffolded Canu assembly was selected for downstream annotation and analysis.

### Annotation

Repeat identification on the selected Canu assembly was performed with RepeatModeler (v.2.0.4), which provided a set of consensus sequences that served as input to RepeatMasker (v.4.1.4) (Table S2) (Flynn et al. 2020). Regions identified as repeats were softmasked in preparation for genome annotation. Six public Illumina RNA-Seq libraries were accessed from NCBI (<u>PRJNA413418</u>) for *J. cinerea* leaf tissue (Table S3). Each library was adapter trimmed with FastP (v.0.23.2) (Chen et al. 2018). The adapter trimmed reads were aligned to the softmasked reference with HISAT2 (v.2.2.1) to provide evidence for gene prediction with an 85% mapping rate cutoff, leading us to exclude two libraries (Kim et al. 2019). The RNA alignments and the softmasked genome served as the input to both BRAKER (v2.1.6) and TSEBRA (v.1.0.1) to generate reference annotations (Bruna et al. 2021; Gabriel et al. 2021). BRAKER was run with transcriptomic alignments (RNA-only mode) and viridiplantae protein input from OrthoDB (v.10) (protein-only mode). Both sets of predictions were fed into TSEBRA. This resulted in one transcriptomic set of predictions from BRAKER and one combined run from TSEBRA. The resulting GFF files were summarized with AGAT (v.1.0.0) and functionally annotated with EnTAP (v.0.10.8) (Dainat 2022; Hart et al. 2020). This provided a reciprocal BLAST estimate against NCBI’s RefSeq Plant Protein database (v208) as well as annotations derived from EggNOG and Gene Ontology (Ashburner et al. 2000; Huerta-Cepas et al. 2019). Both sets of predictions were filtered by the functional annotations; candidates without an EggNOG assignment were removed. The final annotation was selected from a combination of BUSCO, AGAT statistics, and the annotation rate reported by EnTAP.

### Methylation & Circos Visualization

Because CpG and non-CpG methylation are central to plant gene regulation, 5mC methylation was called in all contexts. Bonito-remora (v.0.6.2) with rerio modified base model 5mC_all_context_sup_r1041_e82 was used to estimate methylation (*bonito* n.d.; *rerio* n.d.). The resulting calls were summarized with modbam2bed (v0.9.5) and filtered to >= 10X coverage per site (*modbam2bed*). CpG content was quantified with fastaRegexFinder (v.0.2.0) (Beraldi n.d.). For visualization with ShinyCircos, 5mC methylation was averaged across 100 kb windows, and CpG, repeat, and gene coverage were summed across 100 kb, 10 kb, and 10 kb windows, respectively (Yu et al. 2018).

### Synteny

A macrosynteny analysis was conducted using MCscan (Python version) and jcvi (v.1.3.3) with available *Juglans* species, including *J. cinerea* (PRJEB56451), *J. nigra* (Zhou et al. 2023), *J. regia* (PRJNA291087), and *J. mandshurica* (PRJCA006358) (Tang et al. 2008) (Table 1). If not pre-filtered, any unclassified scaffolds were eliminated from both the genome and annotation files, resulting in a set of 16 chromosomes for each *Juglans* species. Protein-coding sequences from chromosome-scale assemblies, along with their respective annotations (GFF), were used as input for MCscan. The jcvi format module was used to first filter input FASTA files by longest isoform and convert GFF files to BED format. Next, a pairwise synteny search was conducted across *J. cinerea* and *J. mandshurica*, *J. mandshurica* and *J. regia,* and *J. regia* and *J. nigra* respectively. The resulting anchors were then clustered into synteny blocks with a minimum span of 30 and visualized by the jcvi karyotype graphical module. The final chromosome order was determined by length decreasing from left to right. *J. regia* had eight noticeable inversions when compared to *J. mandshurica*; however, upon further inspection these chromosomes were found to be reverse complemented. To address this, the strand orientation in the simple anchors file was manually adjusted.

**Table 1.** Comparison of Juglans chromosome-level genome assemblies and annotations. \**Assembly and annotations not yet publicly available*.

|  | *J. cinerea* | *J. mandshurica* | *J. nigra* | *J. regia* |
| --- | --- | --- | --- | --- |
| **Bioproject** | PRJEB56451 | PRJNA730664 | * | PRJNA291087 |
| **Biosample** | SAMEA111438462 | SAMN19238346 | * | SAMN03704235 |
| **GenBank** | * | GCA_022457165.1 | * | GCA_001411555.2 |
| **Sequence method** | Oxford Nanopore PromethION | PacBio HiFi, Hi-C, Illumina HiSeq | Illumina HiSeq; Pacbio SMRTbell; Hi-C, Illumina NovaSeq6000 | Illumina HiSeq2500; Oxford Nanopore MinION; Hi-C, Illumina HiSeq4000 |
| **Assembly** |  |  |  |  |
| **Genome size (Mb)** | 539.43 | 528.15 | 530.24 | 545.78 |
| **Number of chromosomes** | 16 | 16 | 16 | 16 |
| **N50 (Mb)** | 34.86 | 35.38 | 35.14 | 37.11 |
| **GC content** | 36.64% | 36.49% | 36.36% | 36.08% |
| **Complete BUSCO** | 99.0% | 98.8% | 99.0% | 99.0% |
| **Annotation** |  |  |  |  |
| **Number of genes** | 30,526 | 39,466 | 29,949 | 30,709 |
| **Number of transcripts** | 33,275 | 47,355 | 29,949 | 45,959 |
| **Average gene length (bp)** | 4,187 | 3,748 | 5,175 | 5,309 |
| **Average exon length (bp)** | 209 | 287 | 297 | 291 |
| **Complete BUSCO** | 97.2% | 95.2% | 96.4% | 99.6% |
| **Mono:multi ratio** | 0.207 | 0.347 | 0.210 | 0.206 |
| **Annotation rate** | 98.1% | 84.9% | 96.3% | 99.9% |

### Gene Family Analysis

Orthologous gene families were identified with OrthoFinder (v2.5.4) (Emms and Kelly 2019) (Table S4) using 10 species with reference genomes among *Juglans (J. cinerea, J. nigra, J. regia, J. sigillata, J. microcarpa, J. hindsii, J. mandshurica, J. cathayensis)*, *Carya (Carya illinoinensis)*, and *Pterocarya (Pterocarya stenoptera)*. The species tree, based on resulting gene trees, was constructed in OrthoFinder and rooted using the species tree inference from all genes (STAG) methodology (Emms and Kelly 2018). Support values for internal nodes reflect the proportion of orthogroup trees where the bipartition is observed, and ranges from 0 to 1. Orthogroups only representing one species and orthogroups with oversized representatives from a single species (>100 genes) were removed. The remaining 17,471 orthogroups and ultrametric tree were provided to CAFE v5 (Mendes et al. 2021) to identify gene families with evidence of significant expansion or contraction at p-value < 0.01 using the base model. The longest gene from each orthogroup was used to assign a putative function from EnTAP. Gene families associated with retroelements were filtered from the final analysis.

## Results & Discussion

### Assembly

A total of 59.7 Gb of high accuracy-basecalled (HAC) duplex reads and 59.9 Gb of super-basecalled (SUP) simplex Oxford Nanopore reads were generated using a PromethION sequencer, which each provided 105X coverage at an estimated genome size of 570 Mb (Fig. 1C). Contamination filtering removed primarily bacteria, with the highest proportion of reads associated with *Acinetobacter baumannii* (Fig. S2). Filtering resulted in 58.0 Gb of HAC reads with an N50 length of 19.5 Kb, and 58.0 Gb of SUP reads with an N50 length of 22.4 Kb.

With the SUP reads, Flye initially produced a slightly less complete but more contiguous assembly than Canu; 98.9% vs. 99.2% BUSCO completeness and N50 of 2.55 Mb (1,320 scaffolds) vs. N50 of 2.39 Mb (2,286 contigs), respectively (Table S5; Fig. 1D). Polishing of Flye with Medaka yielded no increase in overall BUSCO completeness, but increased single-copy completeness by 0.1%. Purge Haplotigs increased the contiguity of both assemblies while largely maintaining completeness (N50 of 2.59 Mb in 667 scaffolds for Flye and N50 of 3.02 kb in 345 contigs for Canu). RagTag scaffolding with the *J. mandshurica* assembly drastically increased the contiguity of both assemblies, leaving Flye with 175 scaffolds and Canu with 58 scaffolds. The longest 16 scaffolds from each assembly were extracted and determined to be chromosome-level based on genome size, completeness, and read alignment. *Juglans mandshurica* was chosen for its close phylogenetic position, chromosome conservation, and ability to hybridize (Mu et al. 2017). The assemblies were fairly similar in terms of completeness, contiguity, and overall accuracy (Table S5). The scaffolded Canu assembly was selected for downstream analyses and distinguished only by fewer gaps when compared to the scaffolded Flye assembly (5.32 vs. 9.18 Ns per 100 kb). The final 16 pseudo-chromosome assembly reported is 539 Mb in length with an N50 of 34.86 Mb and an embryophyta BUSCO completeness of 99.0% (Table S5; Fig. 1D). To ensure the assembly was not negatively impacted by the scaffolding process, missassembly was evaluated by examining the percentage of unmapped and soft-clipped bases from read alignment with minimap2. Before scaffolding, 12% of bases were unmapped and 3.58% were soft-clipped. After scaffolding, the percentage of unmapped bases remained the same, and 5.77% of bases were soft-clipped (Fig. S3). This supports minimal impact on overall quality through the scaffolding process.

This exclusively nanopore-assembled genome benefitted from high quality base calls (SUP reads) and >100X coverage to generate a genome of 539 Mb (slightly less than the estimated 570 Mb) (Table S5). The initial genome size estimates were produced from uncorrected long-reads and may not be as reliable of an estimate but were consistent with C-value estimates from Genomes on a Tree (GoaT) (Challis et al. 2023). *Juglans cinerea* is the 10th member of the Juglandaceae to have its whole genome sequenced. Comparing our final *J. cinerea* assembly with that of *J. mandshurica* to which it was scaffolded (Li et al. 2022b), *J. cinerea* exhibits a smaller scaffold N50 size of 34.86 Mb, higher BUSCO completeness score of 99.0%, larger genome size of 539 Mb, and higher GC content of 36.64% (Table 1).

### Annotation

Prior to structural annotation, repeat sequences in the assembly were identified, resulting in 2,353 RepeatScout/RECON families and 1,117 LTRPipeline families. A total of 298 of the LTR families were determined to be redundant and removed, resulting in 3,172 unique repeats. Repeatmasker used this library to softmask 302 Mb (56.02%) of the butternut genome (Fig. 1E). Among the softmasked regions, 28.72% represented retroelements, most of which (22.46%) were predicted to be long terminal repeats (LTRs). Gypsy elements accounted for 8.13% of the genome, whereas Copia elements made up 11.12%. As expected, a large proportion of the repeats (22.42%) were unclassified (Table S2). *J. cinerea* exhibited repeat content that is consistent with other sequenced *Juglans*, with a value that fell between *J. sigillata* (50.06%) and *J. mandshurica* (60.99%) (Ning et al. 2020; Li et al., 2022). Retroelements–mostly LTRs–compose the highest classified repeat content in all three genomes. As reported in the *J. regia* x *Juglans microcarpa* hybrid and *J. regia* genomes, the telomeric repeat CCCTAAA was identified in *J. cinerea* at the ends of each chromosome (Zhu et al. 2019; Marrano et al. 2020).

After trimming, the six RNA-Seq libraries had an average mapping rate of 91.2% against the assembled *J.cinerea* genome (Table S4). Gene prediction was attempted with both BRAKER and TSEBRA on the softmasked assembly (Table S6). BRAKER provided a high quality annotation with only transcriptomic data, resulting in 38,377 predicted genes, 41,357 transcripts, a mono:multi ratio of 0.286, a BUSCO completeness score of 97.2% and an annotation rate of 77.1%. TSEBRA, which combined transcriptomic and protein BRAKER runs, resulted in a slightly lower quality annotation with 35,437 genes, 38,928 transcripts, 0.54 mono:mult ratio, 96.8% BUSCO completeness score, and 86.2% annotation rate. After filtering with EggNOG, the quality of both runs increased, and the number of predicted genes and transcripts was reduced to 30,526 and 33,275 respectively in BRAKER, and 30,328 and 33,694 in TSEBRA. Both programs resulted in a 98.1% annotation rate after filtering, although the BRAKER annotation was closer to the expected mono:multi ratio of plants at 0.21, outperforming that of TSEBRA, 0.36 (Vuruputoor et al. 2022). As a result, BRAKER was chosen to represent the final structural annotation for *J. cinerea*. These annotation results are similar to other *Juglans* species, including the *J. mandshurica* genome released in 2022, which has 39,466 predicted genes, 47,355 transcripts, a mono:multi ratio of 0.347, 95.2% BUSCO completeness, and an annotation rate of 84.9% in chromosome-level scaffolds. Likewise, *J. nigra* and *J. regia*, which have 29,949 and 30,709 genes respectively, 29,949 and 45,959 transcripts, BUSCO completeness of 96.4% and 99.6%, mono:multi of 0.210 and 0.206, and an annotation rate of 96.3% and 99.9% (Table 1). The impressive BUSCO completeness score of the first version of *J. cinerea*, ranking second to version three of *J. regia*, is a testament to the effectiveness of our methodology.

All context methylation (based on 5mC) revealed a central peak pattern to each chromosome that did not correlate with CpG content (Fig. 1E). However, it does appear to correspond with small peaks of repeat content, possibly representing centromeres. Future analyses should analyze methylation by type (CpG, CHG, CHH) in association with repeats and across gene lengths.

### Synteny

The success of our genome assembly is in part due to the quality of the *J. mandshurica* genome, which was used to scaffold our *de novo* assembly. However, utilizing the *scaffold mode* of RagTag and not the *correction mode* allowed some rearrangements to be retained in the *J. cinerea* assembly, as highlighted by the pairwise macrosynteny analysis (Fig. 2). Conserved synteny at chromosome-level was observed in a phylogenetic context, comparing *J. cinerea* to *J. mandshurica*, *J. mandshurica* to *J. nigra,* and *J. nigra* to *J. regia*. A total of 52,993 (NR:38,556) seed synteny blocks (anchors) were found in 294 clusters between *J. cinerea* and *J. mandshurica*, 46,251 (NR:37,316) anchors were identified in 297 clusters between *J.mandshurica* and *J. nigra*, and 47,683 (NR:37,354) anchors were found in 247 cluster across *J.nigra* and *J. regia*. Considering the total number of protein-coding genes in each pairwise comparison, respectively, 68.1%, 66.6%, and 78.6% of compared genomes were collinear, indicating strong support for ancestral collinearity (Li et al. 2022b).

**Figure 2.**
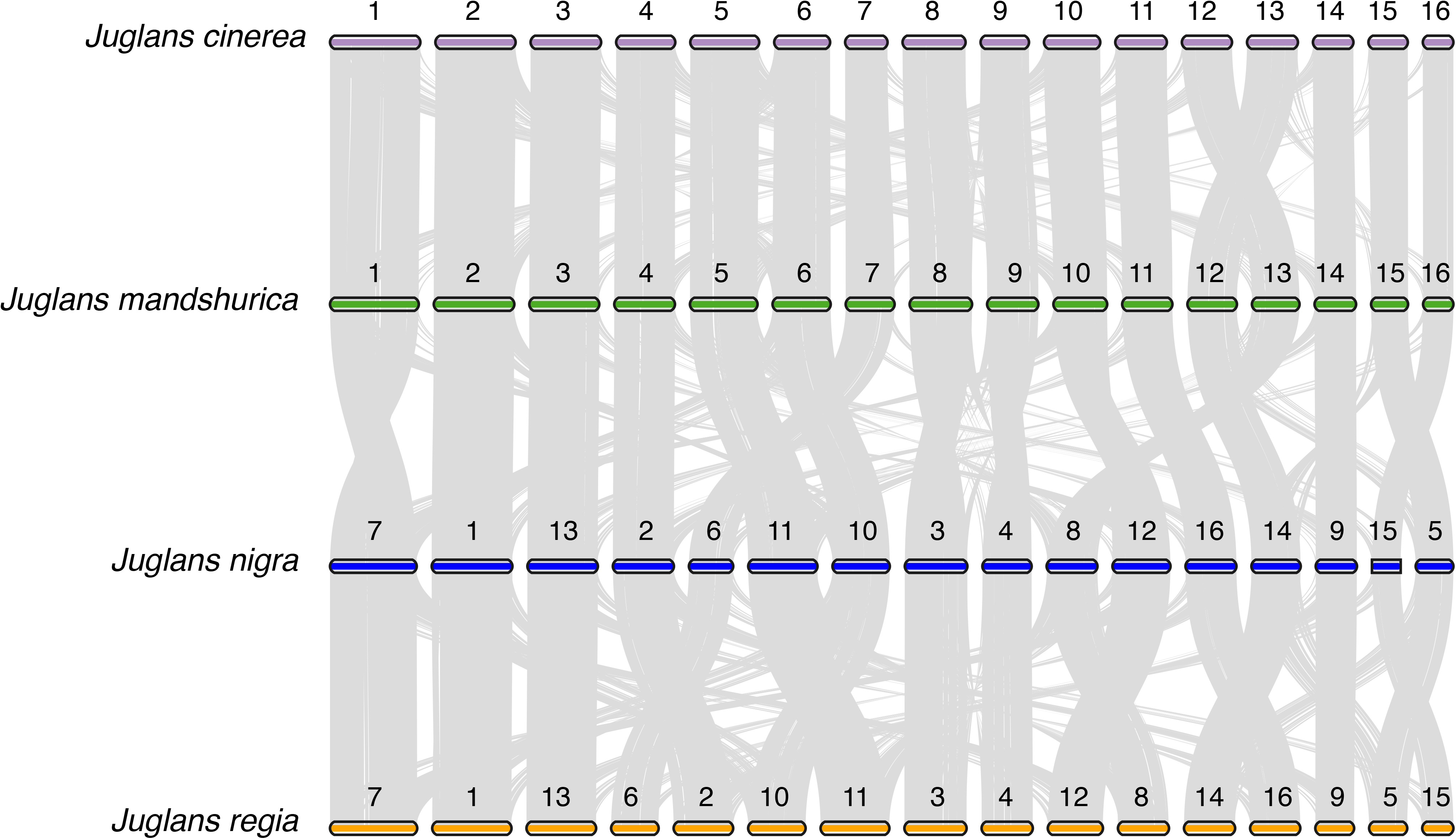
Pairwise macro-synteny analysis in phylogenetic order comparing chromosomes (boxes) and syntenic clusters (grey lines) in J. cinerea (purple) and J. mandshurica (green), J. mandshurica and J. nigra (blue) and J. nigra and J. regia (orange). Published chromosome identifiers and orientation from annotation are maintained, but chromosomes are re-ordered by size (largest to smallest).

### Gene Family Analysis

The OrthoFinder analysis conducted with ten Juglandaceae assigned 384,404 genes out of 401,209 (95.8%) into 33,922 orthogroups (Fig. 3A; Table S4). A total of 13,173 orthogroups (38.8%) were observed in all species and 4,076 orthogroups (12%) were specific to a single species. There were 807 single-copy orthogroups and 16,805 unassigned genes. In *J. cinerea*, 32,743 genes (98.4%) were assigned to an orthogroup. Of the total number of orthgroups found across the 10 species, 22,863 (67.4%) included *J. cinerea*. A total of 132 orthogroups (0.39%) containing 328 genes were unique to *J. cinerea*, and only 532 genes (1.6%) could not be assigned to an orthogroup. *J. cinerea* shares more one-to-one orthologs outside of the *Trachycaryon-Cardiocaryon* clade than within the clade, possibly reflecting isolation and divergence of the ancestral *Cardiocaryon* population to the ancestral *J. cinerea* population (Fig.3A; Table S4).

**Figure 3.**
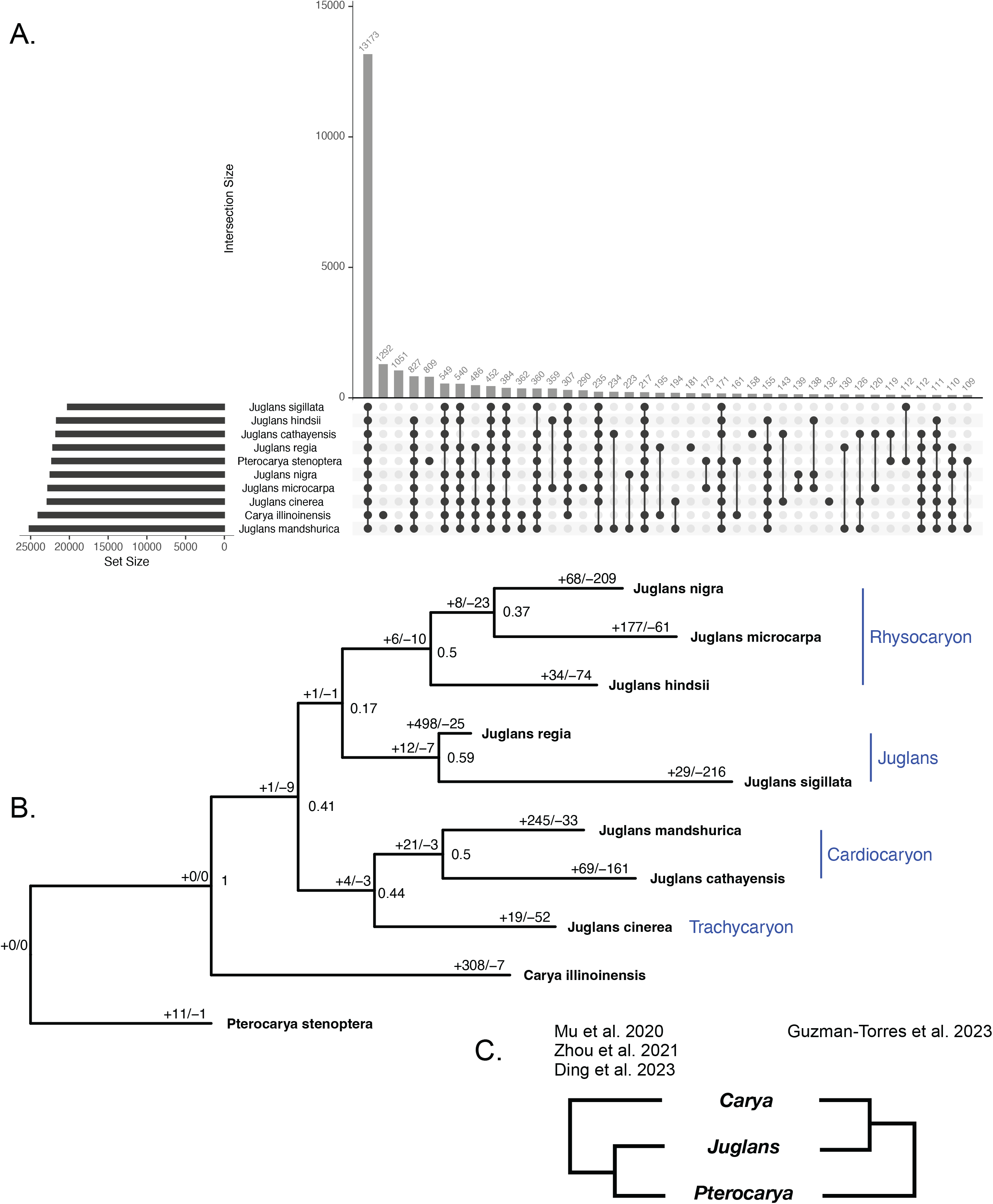
Results of OrthoFinder and the phylogenomic analysis of 33,922 orthrogroup gene trees. Upset plot of the numbers of genes shared by different intersections of 10 species **(A)**. Consensus phylogeny from 33,922 orthogroup gene trees. STAG support values at internal nodes and the number significantly expanding or contracting gene families (p-value < 0.01) on branches **(B)**. Alternate hypotheses of relationships between Carya, Juglans, and Pterocarya using data from RADseq SNPs with de novo alignment (Mu et al. 2020), RADseq SNPs mapped to the J. regia genome (Zhou et al. 2021), subgenomes (Ding et al. 2023) versus whole genome sequencing and orthogroup gene tree analysis (this study) **(C)**.

Members of the Juglandaceae assessed here belong to one of four sections. The North American species *J. nigra*, *J. hindsii* and *J. microcarpa* are members of sect. *Rhysocaryon*. The sampled Eurasian species (sect. *Juglans)* include *J. sigillata and J. regia*, the latter being the predominant cultivated species for walnut production. *Juglans cathayensis and J. mandshurica* are members of sect. *Cardiocaryon*. *J. ailantifolia*, also in section *Cardiocaryon,* but not included here, is sometimes described as a sub-species of *J. mandshurica* (Mu et al. 2017). This is of note given the ability of *J. cinerea* to hybridize with *J. ailantifolia* but not with the sympatric *J. nigra* (Brennan et al. 2020).

Phylogenetic relationships among Juglandaceae remain enigmatic despite multiple sequence and morphology-based analyses. Using protein sequences sourced from whole genome assemblies, *Juglans* was found to be sister to *Carya* with *Pterocarya* as the outgroup (Fig. 3B). Recent phylogenomic analyses relying on sequence data from *de novo* aligned SNPs (Mu et al. 2020), RADseq SNPs mapped to the *J. regia* genome (Zhou et al. 2021), and a deep analysis of subgenomes (Ding et al. 2023) support *Juglans* as sister to *Pterocarya, with Carya* as the outgroup (Fig. 3C). The incongruent results may be an artifact of reduced sampling in this study, however further research is warranted.

Within *Juglans,* phylogenetic discordance is frequently observed between trees built from different datasets, including whole genome sequencing, and the above mentioned SNP sets, subgenome analyses, and plastid versus nuclear markers. The tree generated from this study places *Juglans* section *Rhysocaryon* sister to sect. *Juglans*, in turn, are sister to a clade containing sections *Cadiocaryon* and *Trachycaryon* (Fig. 3C). However, RADseq SNPs dervied from *de novo* alignments generate a tree with section *Juglans* sister to *Cardiocaryon* and section *Rhysocaryon* sister to *Trachycaryon* (Mu et al. 2020), a topology in complete discordance with ours. Topologies made from nuclear SNPs mapped to the *J. regia* genome place *Cardiocaryon* sister to *Trachycaryon*, as observed here; however, section *Juglans* subtends the *Cardiocaryon-Trachycarhon* clade with *Rhysocaryon* as the outgroup in the nuclear SNP-based topology (Zhou et al. 2021).

The placement of *J. cinerea* has varied from the earliest plastid and ITS trees (Stanford et al. 2000), to more recent trees built from whole chloroplast genomes and low-copy nuclear markers (Dong et al. 2017). Characteristic of these trees is the uncertainty of the placement of *J. cinerea* with either Asian or North American *Juglans*. Phylogenetic evidence from whole plastid genomes of *Juglans* indicates *J. cinerea* captured a North American chloroplast genome, while the analysis conducted here places *J. cinerea* sister to the Asian Butternut clade (Fig. 3B). Clearly given the discordance within *Juglans* and *Juglandaceae,* there is a pressing need for additional research using multiple phylogenetic approaches and simulation models.

The overall gene family analysis with CAFE identified a mean gene family birth-death rate (λ) for the whole tree of 0.88, with a tri-modal distribution of gene family expansion/contraction probabilities (Fig. S4). The species with the most rapidly evolving gene families (significant at p-value < 0.01) was *J. regia* with 523 gene families. *J. cinerea* was the species within *Juglans* with the lowest number of significantly evolving gene families, with a total of 71, and the majority of these families were contracting (N = 52, Fig. 3B). After removing orthogroups consisting of retroelements, a total of 54 orthogroups were identified from CAFE as rapidly expanding (9) and rapidly contracting (45) in *J. cinerea* (Fig. 4; Table S7). Among the small set expanding, COG categories broadly described them as cytoskeleton-related, transcription regulation, post-translational modification, cell cycle control, and ion transport. Among those contracting, functions related to transcription regulation, signal transduction, secondary metabolite biosynthesis, carbohydrate transport, disease resistance, and cytoskeleton-related were represented. Examining expanded gene families related to biotic-stress (pathogen/pest resistance), revealed two gene families, one is associated with Auxin response factors (ARFs) and the other with Ripening-related proteins. The pathogenesis-related contracting families were far more numerous and included: NB-LRRs, Wall-associated receptor kinases (WAK), Germin-like proteins, Cytochrome P450s, glycosyl hydrolases, lipolytic acyl hydrolase (LAH), mTERFs, Class III peroxidases (PRX3), UPF0481 proteins, and PAN-like domain proteins. Of these, the following are actually absent in *J. cinerea*: one NB-LRR gene family, one of the three contracting WAK gene families, one cytochrome P450, one of three contracting glycosyl hydrolases gene families, lipolytic acyl hydrolase (LAH), one of two PRX3s, and both families associated with UPF0481 proteins.

**Figure 4.**
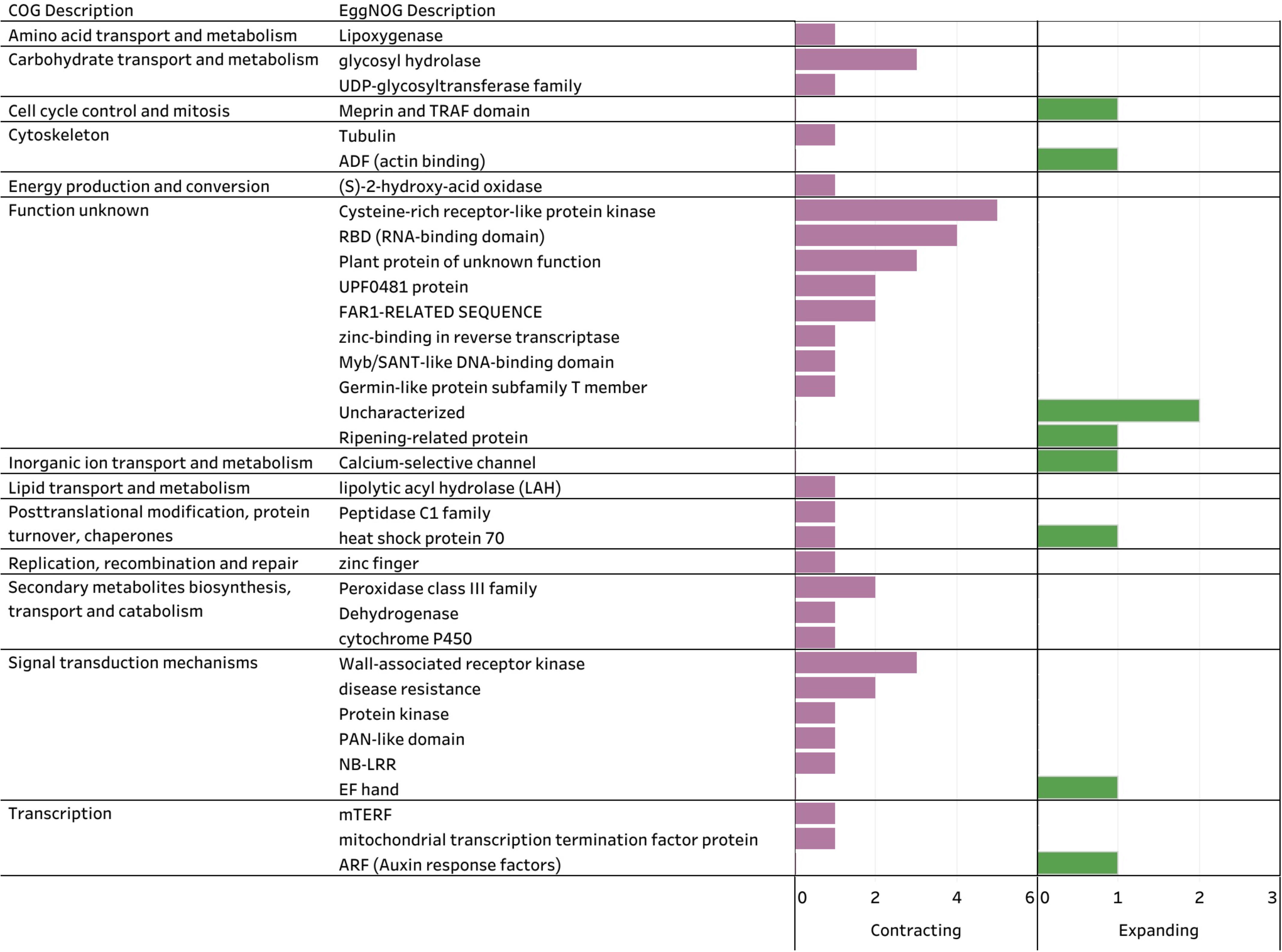
Annotations of J. cinerea significantly contracting (pink) and expanding (green) gene families, excluding retroelement families.

Wall-associated receptor kinases (WAKs) are increasingly associated with plant immunity, including neo- and sub-functionalization in response to changing selection pressure (Stephens et al. 2022; Zhou et al. 2023). Among *Juglans*, they are believed to be absent in *J. hindsii* and *J. sigillata* (Trouern-Trend et al. 2020), rapidly contracting in *J. mandshurica*, and expanding in *J. regia* (Yan et al. 2021). Evaluation of WAK gene families in *J. mandshurica* and *J. regia* found they are under purifying selection (Yan et al. 2021). Expression levels of WAK gene family members were observed to be higher in anthracnose-resistant varieties of *J. regia* (Li et al. 2022a). In *J. cinerea*, a strong signature of contraction across two WAK gene families and absence in the third, was observed. Further investigation of WAKs and other pathogenesis-related families in a population context may reveal more about their role in resistance to specific fungal pathogens.

### Conclusions

The present study reports *Juglans cinerea*’s first high-quality reference genome, generated using Oxford Nanopore technology and a combination of assembly tools. The *J. mandshurica* genome enabled pseudo-chromosome generation and chromosome-level analyses. A preliminary comparative view of *J. cinerea* was presented in context to other species in the Juglandaceae family to assess both synteny and gene family evolution. The construction of the genome was connected to a year-long training program initiated with high molecular weight extraction of the target tree. Students worked closely with graduate students and faculty members to understand the challenges associated with genome assembly and annotation. In doing so, the genome benefitted from robust methods testing. Finally, the program intersected with practitioners who provided input on the reference accession and a framework that will guide the application of this high quality reference in the future. Future studies expanded to population-level analyses of *J. cinerea*, and *J. ailantifolia* hybrid resistance to Oc-j, will be needed to develop conservation strategies informed by genetics.

## Data Availability Statement

The ONT raw reads are available on the European Nucleotide Archive (PRJEB56451). Code, genome assembly, and genome annotation files are available on GitLab: https://doi.org/10.5281/zenodo.5634263.

## Funder Information

Funding was provided by the University of Connecticut, College of Liberal Arts and Sciences, through the Earth and its Future initiative that enabled the initiation of the Biodiversity and Conservation Genomics Training Program. Funding was also provided through National Science Foundation Awards DBI-1943371. Support for nanopore sequencing of the IUCN-listed *Juglans cinerea* was provided by the OrgOne project by Oxford Nanopore Technologies. This research was also funded in part by the USDA Forest Service.The use of trade names is for the information and convenience of the reader and does not imply official endorsement or approval by the U. S. Department of Agriculture or the Forest Service of any product to the exclusion of others that may be suitable.

## Acknowledgements

We thank the Institute for Systems Genomics Center for Genome Innovation at the University of Connecticut for molecular resources and support. We would also like to thank the Computational Biology Core for access to HPC resources. Additional thanks to Patrick G.S. Grady for genome assembly consultation.

## Notes

https://gitlab.com/PlantGenomicsLab/butternut-genome-assembly

https://www.ebi.ac.uk/ena/browser/view/PRJEB56451

### Competing Interest Statement

The authors have declared no competing interest.

### Summary of Updates

Updated language regarding scaffolding with Juglans mandshurica and added related supplemental figure (3). Clarified phylogenetic results. Figure 1A: Re-stylized range map. Figure 2: Reoriented "flipped" chromosomes to clarify syntenic relationships. Figure 3C: Added references.

