## Supplementary figures and images for "Conserving a threatened North American walnut: a chromosome-scale reference genome for butternut (*Juglans cinerea*)"

### Supplemental Figure 1

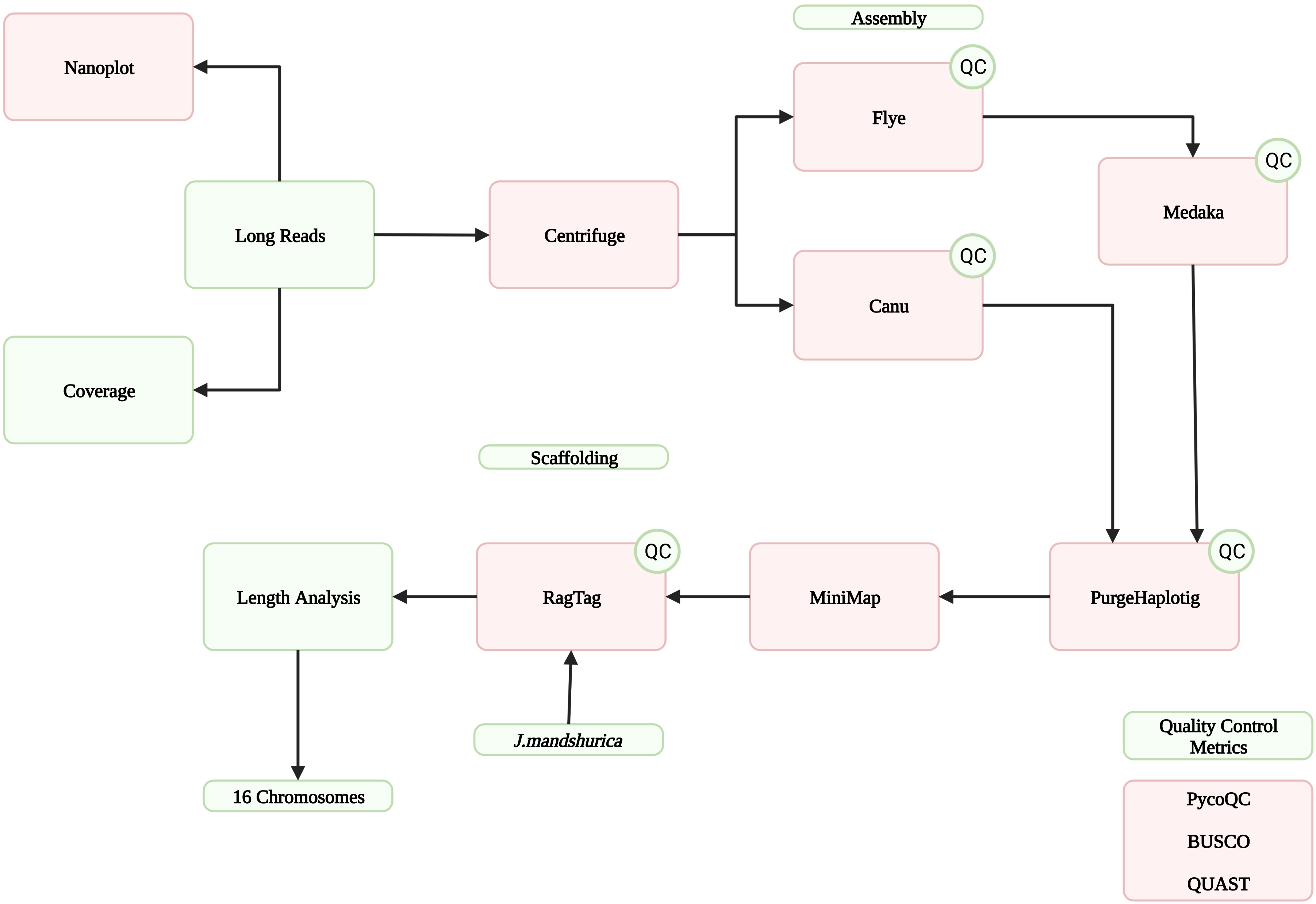

### Supplemental Figure 2

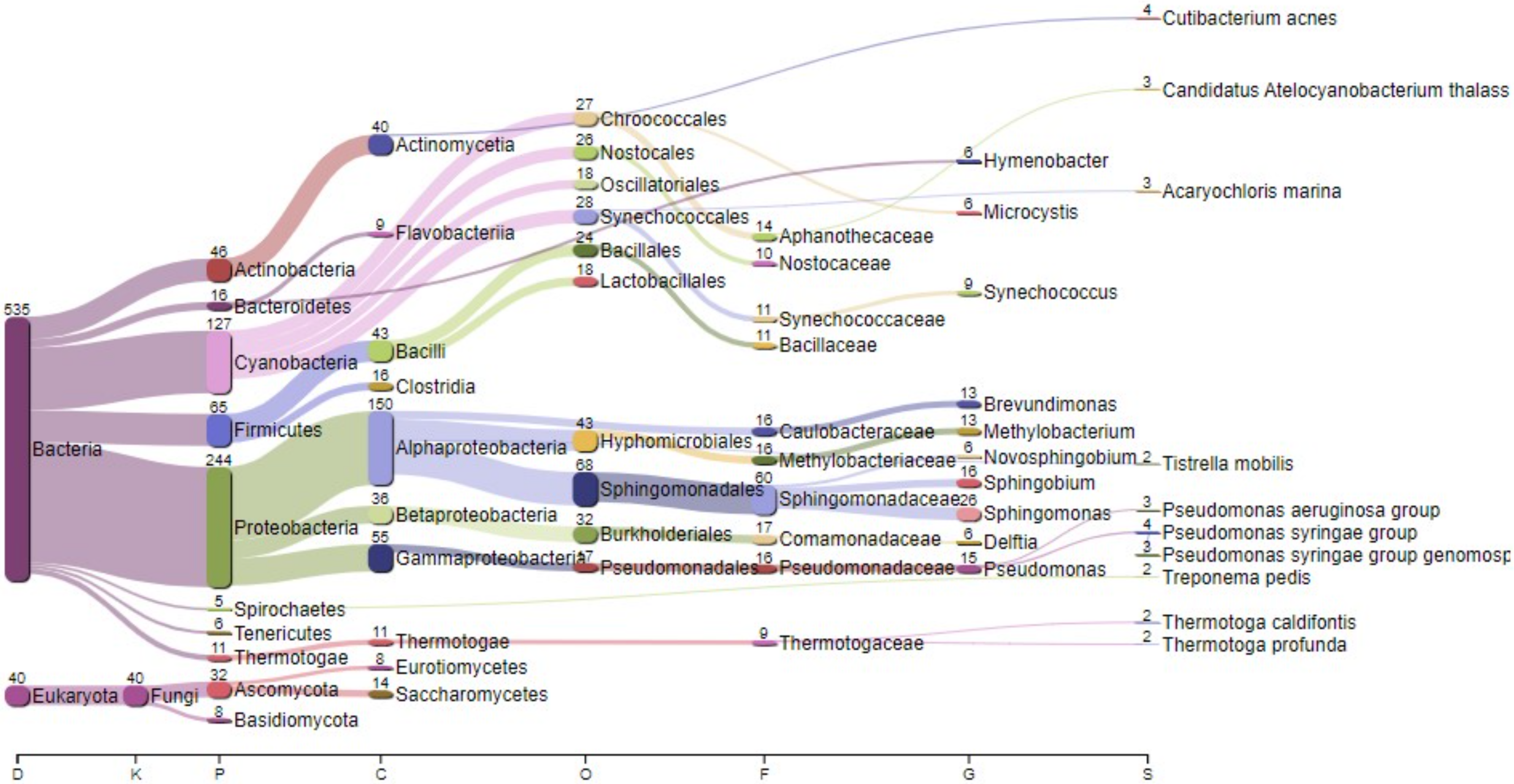

### Supplemental Table 4

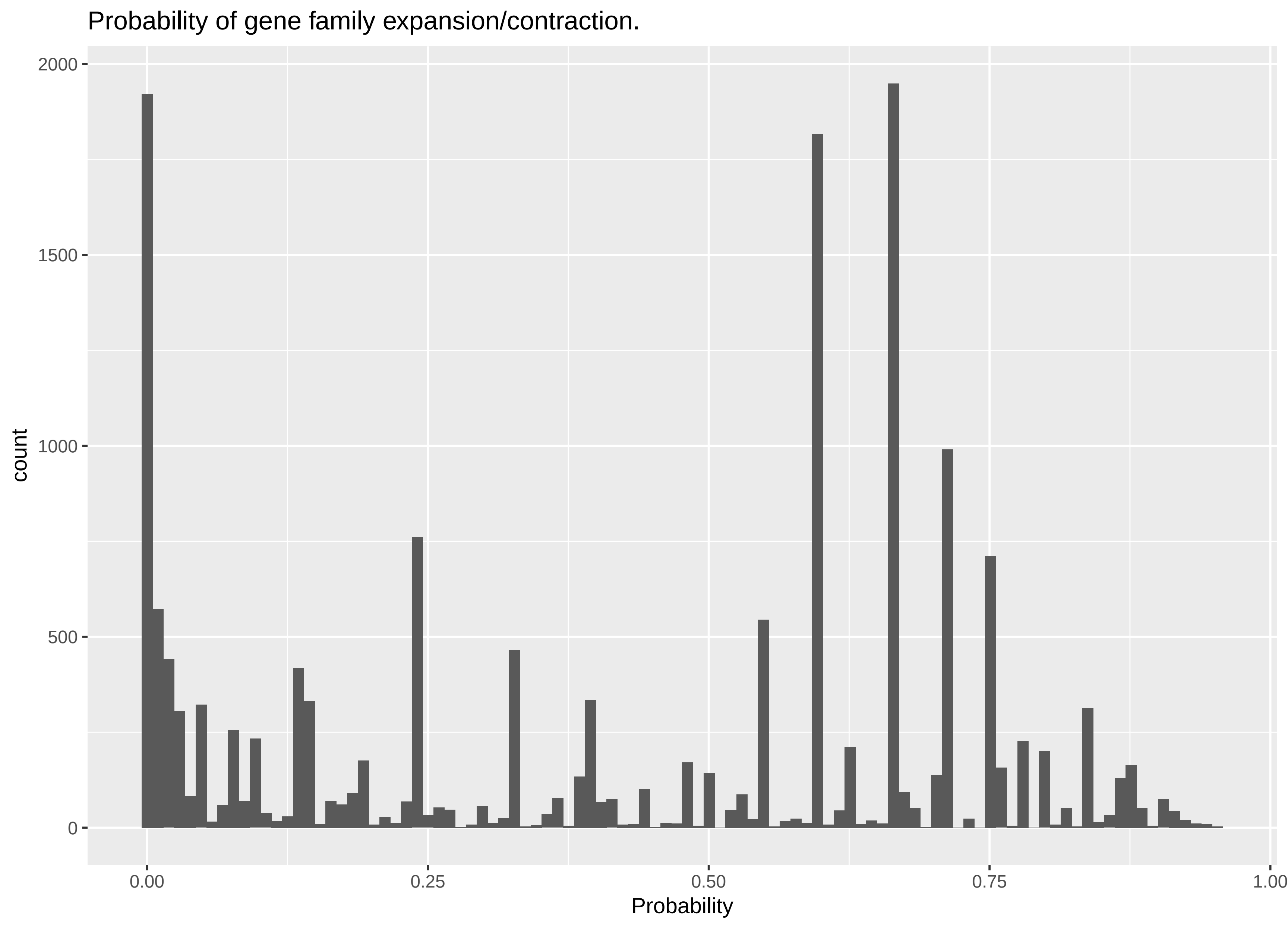
