## Supplemental Figure 3 for "Conserving a threatened North American walnut: a chromosome-scale reference genome for butternut (*Juglans cinerea*)"

**A.** Purged canu assembly

| Bases | Bases Count | % Total | % Aligned |
|---|---|---|---|
| Basecalled | 6.589e+10 | 100% | 100% |
| Unmapped reads | 7.918e+9 | 12.0% | 100% |
| Mapped reads | 5.797e+10 | 88.0% | 100% |
| Softclip | 2.361e+9 | 3.58% | 100% |
| Aligned | 5.561e+10 | 84.4% | 100% |
| Matching | 5.421e+10 | 82.3% | 97.5% |
| Non-matching | 1.404e+9 | 2.13% | 2.52% |
| Insertions | 4.184e+8 | 0.635% | 0.752% |
| Deletions | 4.940e+8 | 0.750% | 0.888% |
| Mismatches | 4.912e+8 | 0.746% | 0.883% |

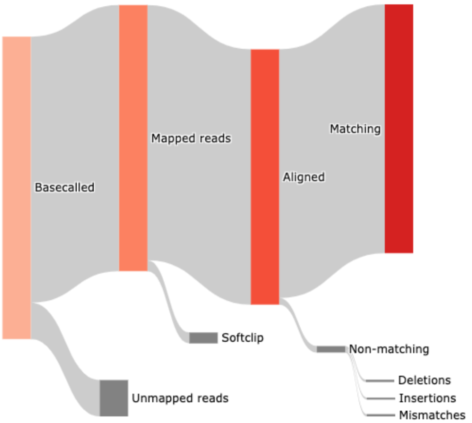


**B.** Scaffolded assembly after purge

| Bases | Bases Count | % Total | % Aligned |
|---|---|---|---|
| Basecalled | 6.589e+10 | 100% | 100% |
| Unmapped reads | 7.929e+9 | 12.0% | 100% |
| Mapped reads | 5.796e+10 | 88.0% | 100% |
| Softclip | 3.801e+9 | 5.77% | 100% |
| Aligned | 5.416e+10 | 82.2% | 100% |
| Matching | 5.250e+10 | 79.7% | 96.9% |
| Non-matching | 1.663e+9 | 2.52% | 3.07% |
| Insertions | 5.136e+8 | 0.780% | 0.948% |
| Deletions | 6.385e+8 | 0.969% | 1.18% |
| Mismatches | 5.111e+8 | 0.776% | 0.944% |

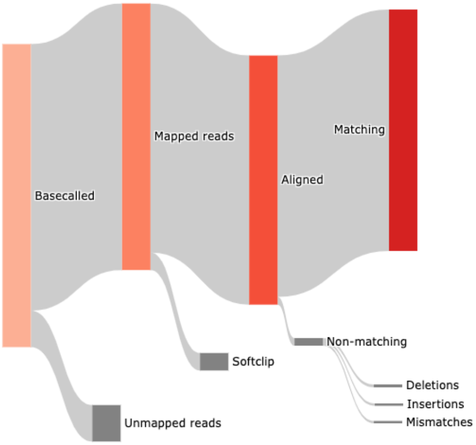


**C.** Pseudo chromosome assembly

| Bases | Bases Count | % Total | % Aligned |
|---|---|---|---|
| Basecalled | 6.589e+10 | 100% | 100% |
| Unmapped reads | 7.962e+9 | 12.1% | 100% |
| Mapped reads | 5.793e+10 | 87.9% | 100% |
| Softclip | 4.865e+9 | 7.38% | 100% |
| Aligned | 5.306e+10 | 80.5% | 100% |
| Matching | 5.137e+10 | 78.0% | 96.8% |
| Non-matching | 1.693e+9 | 2.57% | 3.19% |
| Insertions | 5.150e+8 | 0.782% | 0.970% |
| Deletions | 6.629e+8 | 1.01% | 1.25% |
| Mismatches | 5.150e+8 | 0.782% | 0.971% |

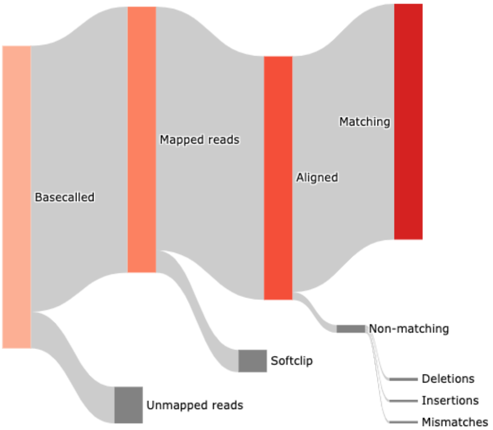
